# *In silico* restriction site analysis of whole genome sequences shows patterns caused by selection and sequence duplications

**DOI:** 10.64898/2026.05.15.725336

**Authors:** Lucia Vedder, Heiko Schoof

**Author notes:** Correspondence: Lucia Vedder.

## Abstract

Biological sequences are known to be not random. Thus, the comparison of *in silico* restriction fragment distributions of random and biological sequences may be an indicator of this non-randomness. Our analyses show that for most of the tested combinations of restriction enzyme and genome sequence the fragments per Megabase of the biological sequence deviate at least more then 10% from the corresponding random sequence. This deviation goes into both directions, i.e. clearly increased values are as common as clearly decreased values. Although there is no species- or restriction-enzyme-specific effect, a clear impact of the GC content both of the restriction site and of the genome sequence can be seen. In contrast to the random sequences, the genome sequences show distinct peaks in their fragment length distributions, hinting to repetitive elements such as transposons.

## Introduction

All biological sequences are affected by natural selection and mutational biases. Thus, they are not entirely random [1]. In biological sequences there are naturally occurring patterns corresponding to functional elements such as e.g. promotors or regulatory elements [2]. In fact, there are also significantly underrepresented sequence patterns, suggesting their absence is due to negative selection [3].

Since the 80s, restriction fragment length polymorphisms are well established as genetic markers [4]. This technique works with the complete digestion of a genome with one or more restriction enzymes. An essential part of these analyses are the restriction sites of the enzyme, which are simple sequence patterns, occurring in biological sequences as well as in random ones.

Given the large numbers of increasingly complete genome sequences, it surprised us that we could not find an analysis of the distribution of restriction sites as an indicator for the non-randomness of biological genome sequences. Therefore, in our paper we want to address the question of how well the distributions of different restriction sites in a small set of representative biological genome sequences match the expectations from corresponding random sequences. Further, we focus our analyses on the possible impact of the GC-content on the frequency of restriction sites.

### *In silico* restriction digest

To analyze the biological impact of a real genomic sequence, a complete *in silico* restriction digest of seven genomes (*Arabidopsis thaliana, Caenorhabditis elegans, Drosophila melanogaster, Daphnia pulex, Mus musculus, Homo sapiens, Thermus aquaticus*) and five random sequences (GC-content of 36%, 40%, 50%, 68%, resp.) was performed. Based on their respective restriction site patterns, 10 restriction enzymes were selected, while taking care of different GC-contents and the possible occurrence of CpG islands. The results of the *in silico* restriction digests are given as the number of fragments per Mb (Megabasepairs, million base pairs) in Tab. 1.

**Tab. 1:** Fragments per Mb of the complete in silico restriction digests of different genomes and random sequences. The sequences and restriction sites are sorted by their respective GC-content. The gray coloring highlights the random sequences. The red cells indicate values at least 1.1x higher than expected based on the corresponding random sequence, the blue cells indicate values at most 10/11x as high as expected based on the corresponding random sequence. The darker coloring of the cells sets these thresholds to more than 1.5x and less than 2/3x, respectively. The bold writing sets these thresholds to more than 2x and less than 1/2x, respectively. C. elegans and A. thaliana are compared to Seq. (A), H. sapiens, D. pulex, M. musculus and D. melanogaster are compared to Seq. (B), Th. aquaticus is compared to Seq. (E).

| Species | GC-content | SspI<br>(AATATT) | AseI<br>(ATTAAT) | EcoRI<br>(GAATTC) | HindIII<br>(AAGCTT) | BamHI<br>(GGATCC) | SacI<br>(GAGCTC) | ApaI<br>(GGGCCC) | SmaI<br>(CCCGGG) | KasI<br>(GGCGCC) | BsePI<br>(GCGCGC) |
| --- | --- | --- | --- | --- | --- | --- | --- | --- | --- | --- | --- |
| *Caenorhabditis elegans* | 35.4% | 1,322.43 | 651.07 | 458.11 | 288.03 | 117.24 | 208.96 | 27.76 | 36.08 | **71.67** | **83.16** |
| *Arabidopsis thaliana* | 36.0% | 992.45 | 770.56 | 309.71 | 560.42 | 145.22 | 173.86 | 23.06 | 26.11 | 28.95 | **9.28** |
| Random sequence (A) | 36.0% | 1,071.27 | 1,072.68 | 339.76 | 340.51 | 105.57 | 107.62 | 33.28 | 34.04 | 34.41 | 34.15 |
| *Homo sapiens* | 39.0% | 760.73 | 468.12 | 272.34 | 276.51 | 119.94 | 198.02 | **154.28** | 126.05 | 77.11 | **24.54** |
| *Daphnia pulex* | 39.5% | 858.27 | 578.17 | 385.23 | 246.72 | 136.85 | 144.06 | 68.69 | 63.28 | 108.43 | 108.11 |
| Random sequence (B) | 40.0% | 733.25 | 735.40 | 325.80 | 325.04 | 143.08 | 142.69 | 63.89 | 63.62 | 62.83 | 65.95 |
| *Mus musculus* | 40.5% | 597.87 | 413.33 | 282.22 | 312.32 | 200.99 | 243.79 | **132.98** | 65.34 | **28.54** | **31.92** |
| *Drosophila melanogaster* | 41.7% | 985.88 | 724.84 | 301.58 | 302.21 | 173.32 | 210.14 | 90.34 | 84.33 | **169.90** | 111.30 |
| Random sequence (C) | 50.0% | 244.38 | 242.11 | 244.99 | 244.81 | 245.58 | 244.43 | 244.79 | 243.73 | 244.98 | 242.69 |
| Random sequence (D) | 50.0% | 243.65 | 243.40 | 244.09 | 243.06 | 245.05 | 246.68 | 242.97 | 245.67 | 244.98 | 245.39 |
| *Thermus aquaticus* | 68.0% | **7.26** | **2.99** | **7.69** | **204.27** | 273.08 | **829.49** | 2,105.13 | 2,452.14 | **508.97** | **125.64** |
| Random sequence (E) | 68.0% | 16.73 | 17.20 | 76.10 | 76.91 | 341.64 | 343.13 | 1,547.25 | 1,552.83 | 1,541.42 | 1,542.52 |

For all sequences a general relation between the GC-content of the sequence and the GC-content of the restriction site can be seen: the closer the GC-content of the sequence and the restriction site, the more fragments per Mb are detected. Thus, lower numbers of fragments per Mb indicate combinations of a high GC-content of the sequence and a low GC-content of the restriction site or vice versa.

The number of fragments per Mb of the random sequences (C) and (D) is in all cases close to 245. This shows that the GC-content of the restriction site has no effect if all bases are equally distributed, e.g. in a random sequence with a CG-content of 50%.

### CpG-islands

To evaluate a possible CpG-island effect, four restriction sites with a GC-content of 100% were selected. Their restriction patterns differ to display different numbers of CpG: the ApaI pattern (GGGCCC) has no CpG, the SmaI (CCCGGG) and the KasI (GGCGCC) pattern have one CpG each and the BsePI pattern (GCGCGC) has two CpGs. In general, a directed effect of the number of CpGs on the number of fragments per Mb (Tab. 1) cannot be seen. Looking only at the two vertebrate species (*H. sapiens* and *M. musculus*) the expected depletion of CpGs in vertebrates [5] is validated by the clearly increased numbers of fragments cut by ApaI (no CpG) and the clearly decreased numbers of fragments cut by BsePI (two CpGs).

### Comparison of random and biological sequences

To compare the biological sequences of the selected species with the computationally created random ones, the number of fragments per Mb of each biological sequence was compared to the respective value of the random sequence with the closest possible GC-content. This means the results of *C. elegans* and *A. thaliana* were compared to random sequence (A), *H. sapiens, D. pulex, M. musculus* and *D. melanogaster* were compared to random sequence (B), and *Th. aquaticus* was compared to random sequence (E).

The results of this comparison are color-marked in Tab. 1. Both higher and lower numbers of fragments than expected are observed in biological sequences, and these appear dependent on the respective combination of species and restriction enzyme with no apparent pattern once the effect of GC-content has been considered. Only fragments cut by AseI (ATTAAT) occur only in lower numbers than expected and only fragments cut by SacI (GAGCTC) occur only in higher numbers than expected.

### Fragment length distributions

The general distribution of restriction fragment lengths can be approximated by a geometric distribution [6], which can be described as a high number of small fragment lengths that decreases exponentially to smaller numbers of high fragment lengths. This general shape can be confirmed by looking at the histograms of the performed *in silico* restriction digests for the analyzed genomes (Fig. 1) as well as for the random sequences (Fig. 2).

**Fig. 1:**
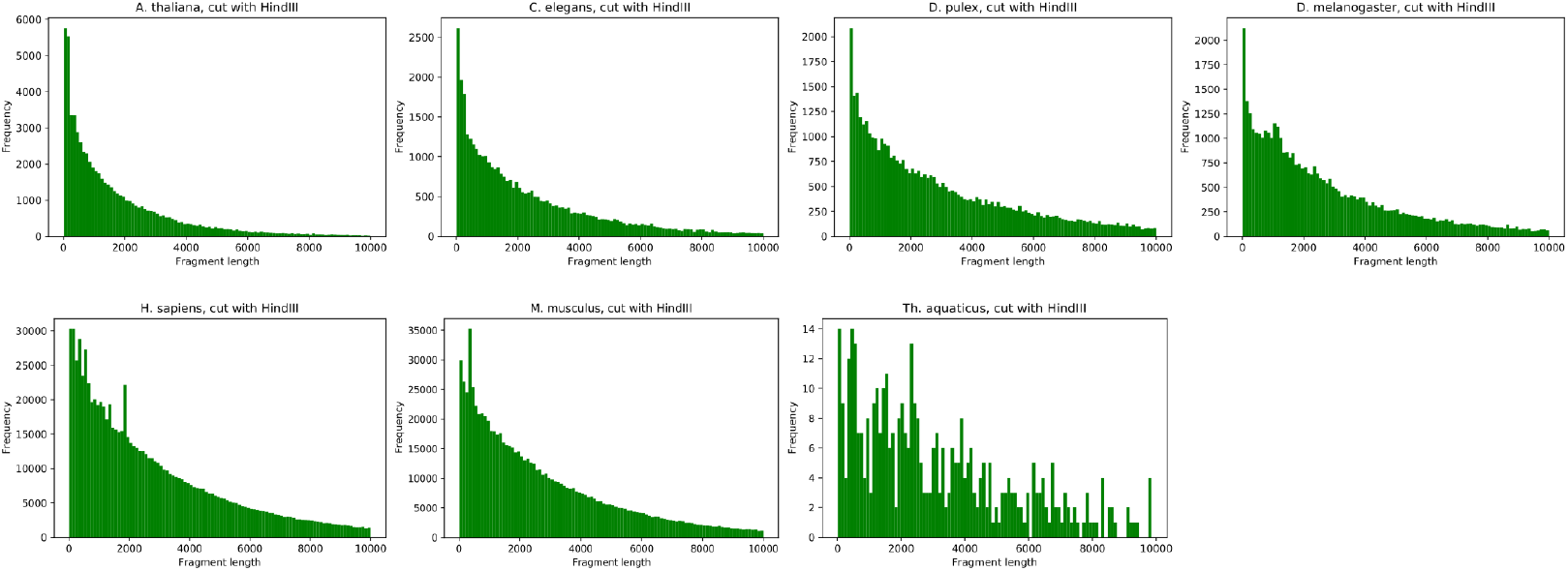
Fragment lengths of the analyzed genomes digested with HindIII. Only fragments up to a length of 10kb are shown.

**Fig. 2:**
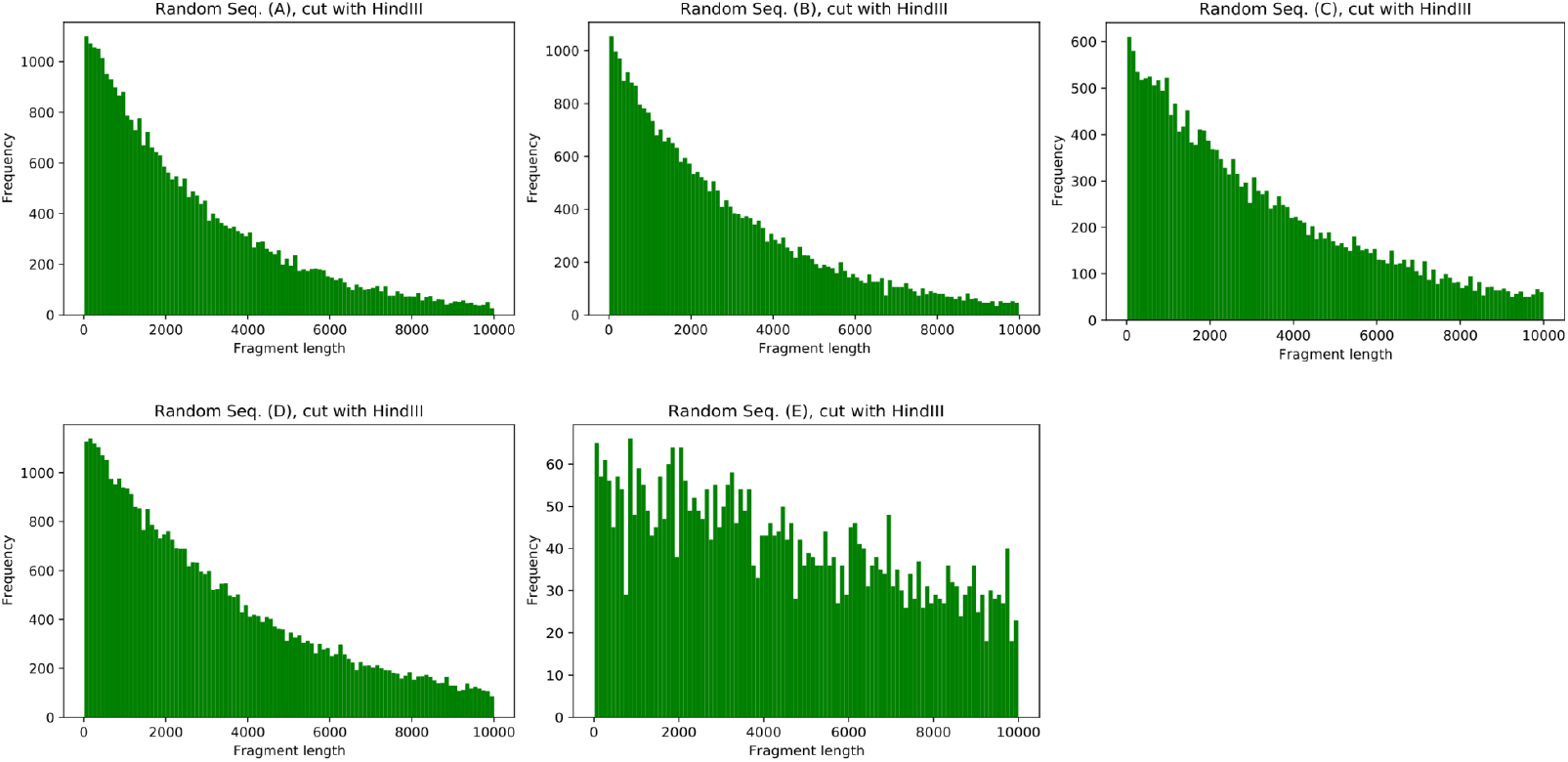
Fragment lengths of the analyzed random sequences digested with HindIII. Only fragments up to a length of 10kb are shown.

Looking at the first 500bp in a higher resolution (Fig. 3) reveals some peaks in the distributions that are not expected from the theoretical distributions as well as from the distributions of the random sequences (Fig. 4). Those peaks are present in almost all analyzed combinations of genomes and restriction sites. They indicate the presence of repetitive elements (e.g. transposons) that contain the respective restriction site, so that a single fragment length is over-represented in the distribution.

**Fig. 3:**
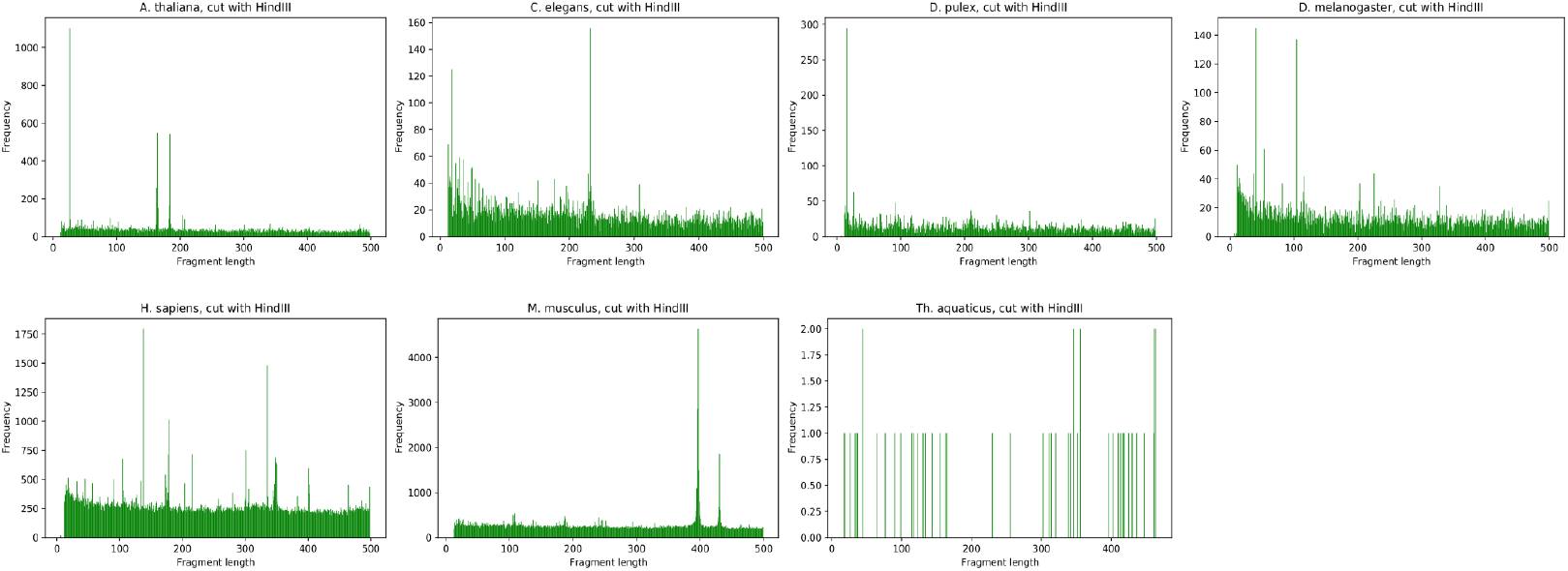
Fragment lengths of the analyzed genomes digested with HindIII. Only fragments up to a length of 500bp are shown.

**Fig. 4:**
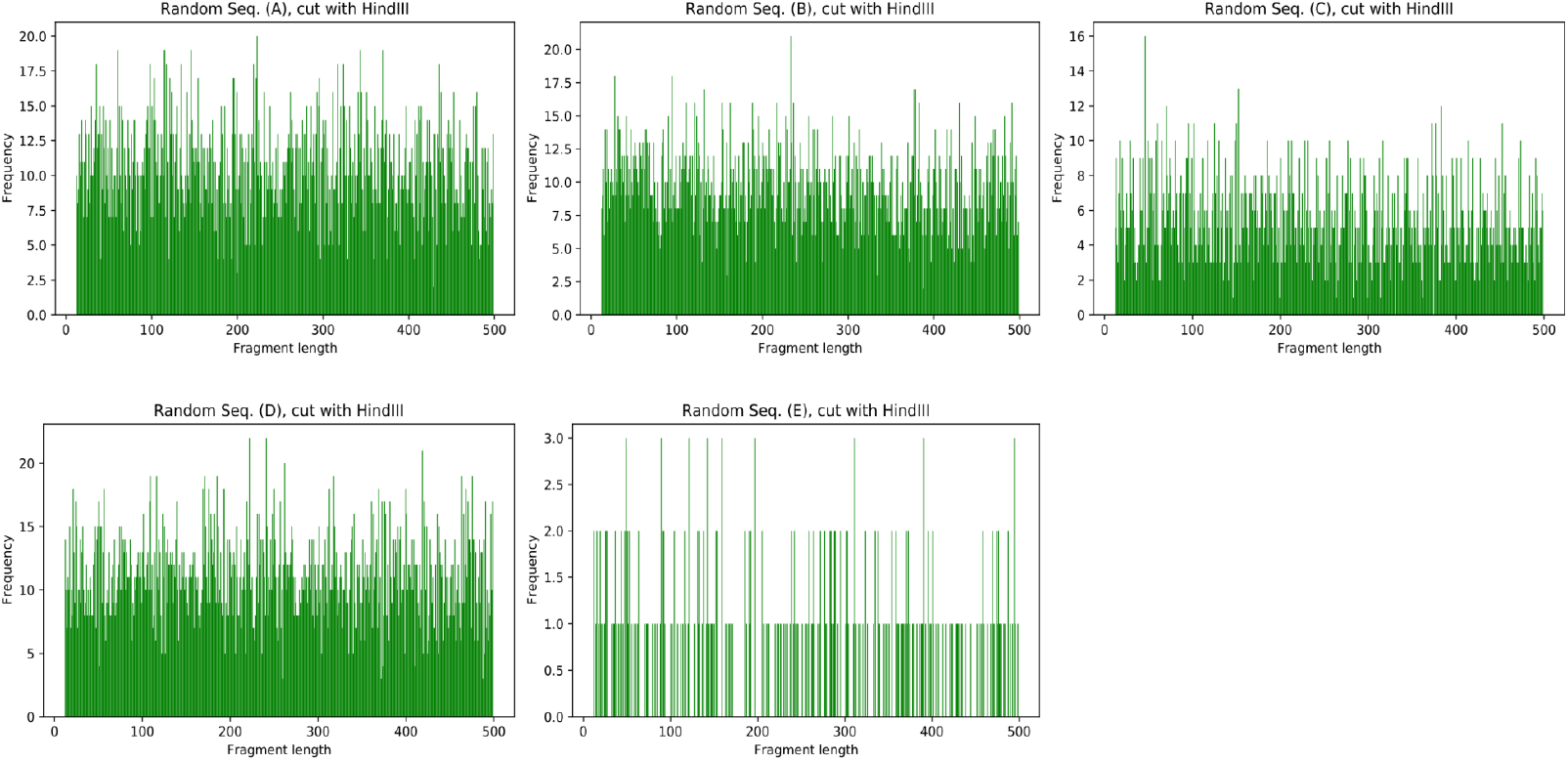
Fragment lengths of the analyzed random sequences digested with HindIII. Only fragments up to a length of 500bp are shown.

We analyzed peaks where the count of a single fragment length deviates more than two standard deviations from the mean of all fragment length counts up to a fragment length of 500bp. Fragments of length <=12bp (fragments that contain only the restriction site) or counts <10 were excluded. These peaks were visually inspected in the respective histogram and only clearly defined peaks relative to the background were retained (Tab. 2). Grouped by species this leaves 16 peaks for *A. thaliana*, 14 for *C. elegans*, 21 for *D. melanogaster*, 24 for *D. pulex*, 32 for *H. sapiens*, 28 for *M. musculus* and 2 for *Th. aquaticus* (Tab. 2).

**Tab. 2:** Selected peaks from the fragment length distributions. All peaks are given as a pair of values, denoted as (fragment length: number of fragments). A peak is defined as a single fragment length, whose count deviates more than two standard deviations from the mean of all fragment length counts up to a fragment length of 500bp. Fragments of length <=12bp (fragments that contain only the restriction site) or counts <10 were excluded. The peaks were selected by visual inspection in the respective histogram. *: No peaks detected. -: No peaks selected.

| Species | Sspl<br>(AATATT) | Asel<br>(ATTAAT) | EcoRI<br>(GAATTC) | HindIII<br>(AAGCTT) | BamHI<br>(GGATCC) | SacI<br>(GAGCTC) | Apal<br>(GGGCCC) | SmaI<br>(CCCGGG) | KasI<br>(GGCGCC) | BsePI<br>(GCGCGC) |
| --- | --- | --- | --- | --- | --- | --- | --- | --- | --- | --- |
| *Arabidopsis thaliana* | - | - | (23: 67)<br>(130: 64)<br>(410: 100) | (26: 1104)<br>(164: 550)<br>(184: 542) | (54: 29)<br>(101: 23)<br>(195: 20)<br>(218: 18)<br>(402: 18) | (36: 41)<br>(78: 24)<br>(181: 25) | - | (254: 11) | (33: 12) | - |
| *Caenorhabditis elegans* | (31: 484)<br>(41: 571)<br>(100: 770) | - | (38: 603) | (18: 125)<br>(232: 156) | (51: 19) | (15: 80)<br>(48: 101)<br>(66: 69)<br>(75: 56) | (15: 58) | (35: 11) | (46: 250) | - |
| *Daphnia pulex* | (179: 403)<br>(211: 271) | - | (18: 132)<br>(68: 95)<br>(168: 67)<br>(409: 99) | (16: 295) | (16:23)<br>(208: 38)<br>(327: 24) | (80: 27)<br>(238: 28)<br>(340: 20) | (20: 21)<br>(39: 33)<br>(103: 20) | (21: 82)<br>(53: 44)<br>(87: 24)<br>(268: 77) | (57: 17)<br>(81: 16)<br>(193: 15)<br>(255: 15) | - |
| *Drosophila melanogaster* | (163: 304) | - | (27: 166)<br>(81: 82)<br>(93: 109)<br>(150: 124)<br>(163: 70)<br>(363: 116) | (41: 145)<br>(54: 61)<br>(104: 129)<br>(226: 44) | (14: 51)<br>(221: 24)<br>(398: 20) | (122: 111)<br>(365: 277) | (119: 79) | (24: 83) | (33: 20)<br>(207: 18)<br>(394: 16) | - |
| *Homo sapiens* | (111: 4793)<br>(315: 7039) | - | (346: 18899) | (138: 1795)<br>(178: 1015)<br>(216: 717)<br>(335: 1479)<br>(348: 687) | (13:256)<br>(18: 264)<br>(45: 178)<br>(69: 168)<br>(142: 181)<br>(192: 149)<br>(242: 159)<br>(416: 128) | (14: 846)<br>(38: 584)<br>(50: 606)<br>(112: 835)<br>(292: 351)<br>(350: 409) | (13: 954)<br>(108: 640)<br>(162: 447) | (29: 695)<br>(38: 908)<br>(56: 1681)<br>(74: 618) | (90: 2421)<br>(133: 439)<br>(182: 342) | - |
| *Mus musculus* | (93: 2892)<br>(115: 2878) | - | (17: 614)<br>(78: 801)<br>(87: 719)<br>(131: 799)<br>(219: 487)<br>(293: 418)<br>(404: 520)<br>(483: 714) | (397: 4637)<br>(431: 1857) | (14: 1305)<br>(18: 1462)<br>(73: 1159)<br>(205: 1273) | (102: 487)<br>(182: 367)<br>(296: 783) | (21: 1680)<br>(205: 2271)<br>(214: 844) | (51: 2355)<br>(58: 3712)<br>(162: 6081) | (70: 243)<br>(89: 957)<br>(129: 671) | - |
| *Thermus aquaticus* | * | * | * | - | - | - | (141: 26)<br>(309: 15) | - | - | - |

For each selected peak, the sequences of all enclosed fragments were clustered with CD-HIT (v4.8.1) [7] using the default parameters. In many cases the resulting number of clusters is quite high, especially for peaks with high sequence numbers. Therefore, only clusters that contain at least 10% of the clustered sequences were analyzed further. In 120 out of 137 peaks there were one or two such clusters (see Tab. S14 in the supplement). The (consensus) sequence of those clusters was searched against the NCBI nr database online using the blastx algorithm (translated nucleotide query against protein database) [8]. Results with descriptions containing ‘hypothetical’, ‘unnamed’, ‘uncharacterized’, ‘putative’, ‘predicted’ and ‘synthetic’ were ignored. This leads to results for at most half of the clusters. Out of 82 results, 30 show matches related to transposable elements, e.g. “gag-pol polyprotein” or “RNA-directed DNA polymerase”. Unlike the other species, sequences of *Homo sapiens* and *Mus musculus* tend to have matches from unrelated species, like e.g. *Plasmodium ovale wallikeri* or *Chlamydia abortus*, respectively. The detailed results are given in Tab. S14 in the supplement. Generally, shorter sequences are more likely to have no database match.

### > 30Mb-fragment in the human genome

In contrast to all other analyzed genomes, the human genome contains one fragment of more than 30Mb for all applied restriction enzyme patterns. The longest fragments of the other genomes are mostly about two orders of magnitude smaller.

Looking at the genomic position of the 30Mb-fragments in different *in silico* digests reveals that they all overlap at the same position on chromosome Y (RefSeq: NC_000024.10). The smallest common region goes from position 26,673,208 to 56,673,220 and is identical with the EcoRI fragment. Unfortunately, there is no annotation available for the complete region within the annotation file of the human genome. At this position the sequence consists of a 30Mb long stretch of N nucleotides, flanked by a EcoRI restriction site. In the chromosome assembly this region is denoted as a gap.

## Discussion

Analyzing our small set of sequences shows that there is clearly an effect of GC-content on the frequency of restriction sites, as shown by the number of fragments per Mb for the given restriction sites with varying GC-content, but other general patterns could not be detected. However, deviations in fragment numbers of at least 10% between the biological genomes and their corresponding random sequences indicate an over-/underrepresentation of the respective restriction site in individual genomes. This supports the idea that biological sequences are not random [1], but are shaped in evolutionary processes. The underrepresentation of AseI sites suggests negative selection [9]. There may be a trend that AT-rich restriction sites are more likely to be underrepresented in species with a high GC-content (such as *Th. aquaticus*) while no systematic overrepresentation of GC-rich sites can be detected. The expected depletion of CpGs in vertebrates [5] is validated within the results by the clearly increased fragment numbers cutting *H. sapiens* and *M. musculus* by ApaI (no CpG) and the clearly decreased fragment numbers cutting the same genomes by BsePI (two CpGs), where both sites have the same GC-content.

Genomes contain sequence repeats of varying sizes [10]. The actual restriction fragment length distributions reveal clearly defined peaks within the genome sequences that are not present in the random sequences. Further analysis of 137 selected peaks shows clear indication that specific fragments are overrepresented. A similarity search of the resulting 199 clusters against the NCBI nr database using BLAST (https://blast.ncbi.nlm.nih.gov/Blast.cgi) [8] reveals meaningful results for 82 (41%) of the clusters. At least 30 out of those 82 clusters match a sequence whose description clearly indicates some kind of transposable element. Thus, the 117 clusters without a meaningful database match may hint at some novel type(s) of transposable element(s).

### Data

The genomes of *Arabidopsis thaliana, Caenorhabditis elegans, Daphnia pulex, Drosophila melanogaster, Homo sapiens, Mus musculus* and *Thermus aquaticus* were downloaded from the NCBI GenomeServer [11]. A list of the respective genome version, genome size, GC-content, number of contigs and N50 is given in the supplement (see Tab. S1).

The random sequences were computationally created using a custom python script (see Methods) in a length of 100Mb each and GC-contents of 36%, 40%, 50% and 68%, respectively. For a GC-content of 50% an additional sequence with a length of 200Mb was created. An overview is given in the supplement (see Tab. S2). The sequence files are published on Zenodo [12].

The restriction enzymes were selected based on the pattern of their restriction site and its respective GC-content. A complete list of the restriction enzymes, the pattern of their restriction site, their GC-content, the number of CpGs (only for restriction sites with 100% GC-content) and their originating species is given in the supplement (see Tab. S3).

## Methods

All analyses were made *in silico* using custom Python (v.3.8.5) [13] scripts. The scripts are provided on GitHub (https://github.com/lu-vedder/RestrictionEnzymeFragments) and published on Zenodo [14].

The random sequences were computed using the python script random_sequence.py. The first analysis step was the fragmentation of the given genomes at the respective restriction site (`fragment_genome.py`). Each fragment starts with either the first base of its contig or the complete restriction site pattern and ends with either the last base of its contig or the complete restriction site pattern. Thus, the restriction site pattern is doubled at each *in silico* cut. Therefore, the minimal possible fragment length for fragments starting/ending at the contig’s first/last base, respectively, is 6 (the length of the pattern). The minimal possible fragment length for fragments whose both sides are within the contig is 12 (the length of two directly following patterns).

Some basic statistics of the length distributions of the fragments are computed in a second step (`fragment_statistics.py`). The important number of fragments per Mb is calculated as the total number of fragments divided by the size of the genome in Mb. The complete statistics are given in the supplement (see Tab. S4-S13).

The histograms shown in Fig. 1-4 are created using the plot_fragments.py script.

The analyzed peaks are detected using the find_peaks.py script.

The statistical values on the analyzed genomes in Tab. S1 were computed using `genome_statistics.py`.

## Acknowledgments

Alex Olek for inspiring a comprehensive survey and for discussions. Ark Biodiversity GmbH for financial support.

## Supporting information captions

**Tab. S1:** Genomes, used for the *in silico* fragmentation. The genomes were downloaded from the NCBI GenomeServer [11]. Statistical values were computed using a custom python script [14].

**Tab. S2:** Random sequences, used for the *in silico* fragmentation. The sequences are published on Zenodo [12]. The custom python script used for generating the random sequences is also published on Zenodo [14].

**Tab. S3:** Restriction enzymes, used for *in silico* fragmentation. The enzymes were selected based on the pattern of their respective restriction site using EnzymeFinder (v2.8.1, http://enzymefinder.neb.com). The originating species of the restriction enzymes were looked up in the REBASE database (http://rebase.neb.com).

**Tab. S4:** Fragment statistics for the *in silico* fragmentation with SspI.

**Tab. S5:** Fragment statistics for the *in silico* fragmentation with AseI.

**Tab. S6:** Fragment statistics for the *in silico* fragmentation with EcoRI.

**Tab. S7:** Fragment statistics for the *in silico* fragmentation with HindIII.

**Tab. S8:** Fragment statistics for the *in silico* fragmentation with BamHI.

**Tab. S9:** Fragment statistics for the *in silico* fragmentation with SacI.

**Tab. S10:** Fragment statistics for the *in silico* fragmentation with ApaI.

**Tab. S11:** Fragment statistics for the *in silico* fragmentation with SmaI.

**Tab. S12:** Fragment statistics for the *in silico* fragmentation with KasI.

**Tab. S13:** Fragment statistics for the *in silico* fragmentation with BsePI.

**Tab. S14:** Complete list of the clusters of the selected peaks.

