## Supplement Tables S1 - S13 for "*In silico* restriction site analysis of whole genome sequences shows patterns caused by selection and sequence duplications"

***Tab. S1:*** *Genomes, used for the in silico fragmentation. The genomes were downloaded from the NCBI GenomeServer [11]. Statistical values were computed using a custom python script [14].*

| Species | Genome version | Genome size (Mb) | GC-content | No. contigs | N50 (Mb) |
|---|---|---|---|---|---|
| *Arabidopsis thaliana* | GCF_000001735.4_TAIR10.1_genomic.fna | 119.67 | 36.0% | 7 | 23.46 |
| *Caenorhabditis elegans* | GCF_000002985.6_WBcel235_genomic.fna | 100.29 | 35.4% | 7 | 17.49 |
| *Daphnia pulex* | GCA_900092285.2_PA42_4.1_genomic.fna | 189.55 | 39.5% | 493 | 1.16 |
| *Drosophila melanogaster* | GCF_000001215.4_Release_6_plus_ISO1_MT_genomic.fna | 143.73 | 41.7% | 1,870 | 25.29 |
| *Homo sapiens* | GCF_000001405.39_GRCh38.p13_genomic.fna | 3,272.09 | 39.0% | 639 | 145.14 |
| *Mus musculus* | GCF_000001635.27_GRCm39_genomic.fna | 2,728.22 | 40.5% | 61 | 130.53 |
| *Thermus aquaticus* | GCF_001399775.1_ASM139977v1_genomic.fna | 2.34 | 68.0% | 5 | 2.16 |

***Tab. S2:*** *Random sequences, used for the in silico fragmentation. The sequences are published on Zenodo [12]. The custom python script used for generating the random sequences is also published on Zenodo [14].*

| Name | Size (Mb) | GC-content |
|---|---|---|
| Random sequence (A) | 100.0 | 36.0% |
| Random sequence (B) | 100.0 | 40.0% |
| Random sequence (C) | 100.0 | 50.0% |
| Random sequence (D) | 200.0 | 50.0% |
| Random sequence (E) | 100.0 | 68.0% |

***Tab. S3:*** *Restriction enzymes, used for in silico fragmentation. The enzymes were selected based on the pattern of their respective restriction site using EnzymeFinder (v2.8.1, http://enzymefinder.neb.com). The originating species of the restriction enzymes were looked up in the REBASE database (http://rebase.neb.com).*

| **Restriction enzyme** | **Pattern** | **GC-content** | **No. CpGs** | **Originating species** |
|---|---|---|---|---|
| SspI | AATATT | 0.0% | - | *Sphaerotilus species* |
| AseI | ATTAAT | 0.0% | - | *Aquaspirillum serpens* |
| EcoRI | GAATTC | 33.3% | - | *Escherichia coli* RY13 |
| HindIII | AAGCTT | 33.3% | - | *Haemophilus influenzae* Rd |
| BamHI | GGATCC | 66.7% | - | *Bacillus amyloliquefaciens* H |
| SacI | GAGCTC | 66.7% | - | *Streptomyces achromogenes* |
| ApaI | GGGCCC | 100.0% | 0 | *Acetobacter pasteurianus* sub. pasteurianus |
| SmaI | CCCGGG | 100.0% | 1 | *Serratia marcescens* Sb |
| KasI | GGCGCC | 100.0% | 1 | *Kluyvera ascorbata* |
| BsePI | GCGCGC | 100.0% | 2 | *Bacillus stearothermophilus* P6 |

***Tab. S4:*** *Fragment statistics for the in silico fragmentation with SspI.*

| **Species** | **No. fragments** | **Min. fragment length (bp)** | **Max. fragment length (bp)** | **Most freq. fragment length (bp)** | **Fragments per Mb** |
|---|---|---|---|---|---|
| *Arabidopsis thaliana* | 118,766 | 12 | 68,490 | 38 | 992.45 |
| *Caenorhabditis elegans* | 132,627 | 12 | 50,213 | 100 | 1,322.43 |
| *Daphnia pulex* | 162,685 | 8 | 182,524 | 179 | 858.27 |
| *Drosophila melanogaster* | 141,701 | 7 | 56,467 | 14 | 985.88 |
| *Homo sapiens* | 2,489,162 | 8 | 30,155,042 | 315 | 760.73 |
| *Mus musculus* | 1,631,129 | 12 | 3,062,340 | 1,043 | 597.87 |
| *Thermus aquaticus* | 17 | 195 | 350,363 | - | 7.26 |
| Random sequence (A) | 107,127 | 12 | 10,234 | 48 | 1,071.27 |
| Random sequence (B) | 73,325 | 12 | 14,555 | 26 | 733.25 |
| Random sequence (C) | 24,438 | 12 | 47,478 | 194 | 244.38 |
| Random sequence (D) | 48,729 | 12 | 43,617 | 199 | 243.65 |
| Random sequence (E) | 1,673 | 28 | 458,982 | 3,986 | 16.73 |

***Tab. S5:*** *Fragment statistics for the in silico fragmentation with AseI.*

| **Species** | **No. fragments** | **Min. fragment length (bp)** | **Max. fragment length (bp)** | **Most freq. fragment length (bp)** | **Fragments per Mb** |
|---|---|---|---|---|---|
| *Arabidopsis thaliana* | 92,213 | 10 | 78,767 | 10 | 770.56 |
| *Caenorhabditis elegans* | 65,296 | 10 | 39,066 | 10 | 651.07 |
| *Daphnia pulex* | 109,592 | 9 | 74,283 | 10 | 578.17 |
| *Drosophila melanogaster* | 104,181 | 6 | 78,868 | 10 | 724.84 |
| *Homo sapiens* | 1,531,718 | 10 | 30,039,106 | 10 | 468.12 |
| *Mus musculus* | 1,127,663 | 10 | 3,061,944 | 10 | 413.33 |
| *Thermus aquaticus* | 7 | 3,236 | 1,278,573 | - | 2.99 |
| Random sequence (A) | 107,268 | 10 | 12,430 | 10 | 1,072.68 |
| Random sequence (B) | 73,540 | 10 | 15,485 | 10 | 735.40 |
| Random sequence (C) | 24,211 | 10 | 45,439 | 10 | 242.11 |
| Random sequence (D) | 48,680 | 10 | 41,790 | 10 | 243.40 |
| Random sequence (E) | 1,720 | 10 | 411,546 | 14,681 | 17.20 |

***Tab. S6:*** *Fragment statistics for the in silico fragmentation with EcoRI.*

| Species | No. fragments | Min. fragment length (bp) | Max. fragment length (bp) | Most freq. fragment length (bp) | Fragments per Mb |
|---|---|---|---|---|---|
| *Arabidopsis thaliana* | 37,063 | 12 | 72,592 | 410 | 309.71 |
| *Caenorhabditis elegans* | 45,944 | 12 | 54,431 | 38 | 458.11 |
| *Daphnia pulex* | 73,021 | 7 | 74,786 | 18 | 385.23 |
| *Drosophila melanogaster* | 43,346 | 6 | 138,018 | 27 | 301.58 |
| *Homo sapiens* | 891,106 | 6 | 30,000,012 | 346 | 272.34 |
| *Mus musculus* | 769,971 | 6 | 3,070,023 | 1,379 | 282.22 |
| *Thermus aquaticus* | 18 | 695 | 575,486 | - | 7.69 |
| Random sequence (A) | 33,976 | 12 | 33,922 | 159 | 339.76 |
| Random sequence (B) | 32,580 | 12 | 36,218 | 182 | 325.80 |
| Random sequence (C) | 24,499 | 12 | 40,224 | 91 | 244.99 |
| Random sequence (D) | 48,817 | 12 | 44,773 | 163 | 244.09 |
| Random sequnece (E) | 7,610 | 12 | 121,502 | 2,428 | 76.10 |

***Tab. S7:*** *Fragment statistics for the in silico fragmentation with HindIII.*

| Species | No. fragments | Min. fragment length (bp) | Max. fragment length (bp) | Most freq. fragment length (bp) | Fragments per Mb |
|---|---|---|---|---|---|
| *Arabidopsis thaliana* | 67,065 | 12 | 60,164 | 26 | 560.42 |
| *Caenorhabditis elegans* | 38,916 | 12 | 52,852 | 232 | 288.03 |
| *Daphnia pulex* | 46,765 | 6 | 96,854 | 16 | 246.72 |
| *Drosophila melanogaster* | 43,436 | 6 | 136,725 | 41 | 302.21 |
| *Homo sapiens* | 904,778 | 6 | 30,216,696 | 138 | 276.51 |
| *Mus musculus* | 852,077 | 12 | 3,070,057 | 397 | 312.32 |
| *Thermus aquaticus* | 478 | 17 | 39,963 | 1,710 | 204.27 |
| Random sequence (A) | 34,051 | 12 | 35,332 | 523 | 340.51 |
| Random sequence (B) | 32,504 | 12 | 33,710 | 233 | 325.04 |
| Random sequence (C) | 24,481 | 12 | 43,552 | 46 | 244.81 |
| Random sequence (D) | 48,612 | 12 | 48,483 | 663 | 243.06 |
| Random sequence (E) | 7,691 | 12 | 116,933 | 5,146 | 76.91 |

***Tab. S8:** Fragment statistics for the in silico fragmentation with BamHI.*

| **Species** | **No. fragments** | **Min. fragment length (bp)** | **Max. fragment length (bp)** | **Most freq. fragment length (bp)** | **Fragments per Mb** |
|---|---|---|---|---|---|
| *Arabidopsis thaliana* | 17,378 | 12 | 138,871 | 12 | 145.22 |
| *Caenorhabditis elegans* | 11,758 | 12 | 131,094 | 51 | 117.24 |
| *Daphnia pulex* | 25,940 | 12 | 122,341 | 208 | 136.85 |
| *Drosophila melanogaster* | 24,911 | 7 | 136,502 | 1,317 | 173.32 |
| *Homo sapiens* | 392,460 | 7 | 30,119,691 | 2,063 | 119.94 |
| *Mus musculus* | 548,353 | 7 | 3,090,732 | 514 | 200.99 |
| *Thermus aquaticus* | 639 | 15 | 26,504 | 73 | 273.08 |
| Random sequence (A) | 10,557 | 12 | 96,148 | 1,389 | 105.57 |
| Random sequence (B) | 14,308 | 12 | 67,313 | 350 | 143.08 |
| Random sequence (C) | 24,558 | 12 | 37,480 | 154 | 245.58 |
| Random sequence (D) | 49,010 | 12 | 44,576 | 329 | 245.05 |
| Random sequence (E) | 34,164 | 12 | 42,858 | 151 | 341.64 |

***Tab. S9:** Fragment statistics for the in silico fragmentation with SacI.*

| **Species** | **No. fragments** | **Min. fragment length (bp)** | **Max. fragment length (bp)** | **Most freq. fragment length (bp)** | **Fragments per Mb** |
|---|---|---|---|---|---|
| *Arabidopsis thaliana* | 20,806 | 12 | 88,261 | 512 | 173.86 |
| *Caenorhabditis elegans* | 20,957 | 12 | 74,368 | 48 | 208.96 |
| *Daphnia pulex* | 27,307 | 12 | 124,612 | 238 | 144.06 |
| *Drosophila melanogaster* | 30,203 | 7 | 192,434 | 365 | 210.14 |
| *Homo sapiens* | 647,949 | 6 | 30,221,429 | 14 | 198.02 |
| *Mus musculus* | 665,120 | 12 | 3,077,435 | 296 | 243.79 |
| *Thermus aquaticus* | 1,941 | 15 | 9,936 | 30 | 829.49 |
| Random sequence (A) | 10,762 | 12 | 105,992 | 1,328 | 107.62 |
| Random sequence (B) | 14,269 | 14 | 72,172 | 258 | 142.69 |
| Random sequence (C) | 24,443 | 12 | 46,284 | 265 | 244.43 |
| Random sequence (D) | 49,336 | 12 | 48,223 | 232 | 246.68 |
| Random sequence (E) | 34,313 | 12 | 29,333 | 214 | 343.13 |

***Tab. S10:*** *Fragment statistics for the in silico fragmentation with ApaI.*

| Species | No. fragments | Min. fragment length (bp) | Max. fragment length (bp) | Most freq. fragment length (bp) | Fragments per Mb |
|---|---|---|---|---|---|
| *Arabidopsis thaliana* | 2,760 | 14 | 538,805 | 25 | 23.06 |
| *Caenorhabditis elegans* | 2,784 | 15 | 353,101 | 15 | 27.76 |
| *Daphnia pulex* | 13,021 | 8 | 212,560 | 39 | 68.69 |
| *Drosophila melanogaster* | 12,985 | 8 | 186,801 | 119 | 90.34 |
| *Homo sapiens* | 504,807 | 12 | 30,233,887 | 13 | 154.28 |
| *Mus musculus* | 362,790 | 12 | 3,103,913 | 205 | 132.98 |
| *Thermus aquaticus* | 4,926 | 12 | 6,093 | 141 | 2,105.13 |
| Random sequence (A) | 3,328 | 17 | 264,528 | 2,129 | 33.28 |
| Random sequence (B) | 6,389 | 13 | 143,695 | 6,937 | 63.89 |
| Random sequence (C) | 24,479 | 12 | 50,794 | 302 | 244.79 |
| Random sequence (D) | 48,594 | 12 | 42,397 | 683 | 242.97 |
| Random sequence (E) | 154,725 | 12 | 7,427 | 14 | 1,547.25 |

***Tab. S11:*** *Fragment statistics for the in silico fragmentation with SmaI.*

| Species | No. fragments | Min. fragment length (bp) | Max. fragment length (bp) | Most freq. fragment length (bp) | Fragments per Mb |
|---|---|---|---|---|---|
| *Arabidopsis thaliana* | 3,125 | 10 | 474,214 | 504 | 26.11 |
| *Caenorhabditis elegans* | 3,618 | 12 | 329,548 | 35 | 36.08 |
| *Daphnia pulex* | 11,994 | 12 | 297,650 | 21 | 63.28 |
| *Drosophila melanogaster* | 12,121 | 12 | 217,110 | 24 | 84.33 |
| *Homo sapiens* | 412,447 | 11 | 30,207,030 | 56 | 126.05 |
| *Mus musculus* | 178,256 | 12 | 3,153,549 | 162 | 65.34 |
| *Thermus aquaticus* | 5,738 | 12 | 5,439 | 12 | 2,452.14 |
| Random sequence (A) | 3,404 | 19 | 318,997 | 14,123 | 34.04 |
| Random sequence (B) | 6,362 | 12 | 143,178 | 3,283 | 63.62 |
| Random sequence (C) | 24,373 | 12 | 46,704 | 438 | 243.73 |
| Random sequence (D) | 49,134 | 12 | 44,717 | 89 | 245.67 |
| Random sequence (E) | 155,283 | 12 | 9,644 | 13 | 1,552.83 |

***Tab. S12:*** *Fragment statistics for the in silico fragmentation with KasI.*

| Species | No. fragments | Min. fragment length (bp) | Max. fragment length (bp) | Most freq. fragment length (bp) | Fragments per Mb |
|---|---|---|---|---|---|
| *Arabidopsis thaliana* | 3,464 | 12 | 331,150 | 33 | 28.95 |
| *Caenorhabditis elegans* | 7,188 | 12 | 165,005 | 46 | 71.67 |
| *Daphnia pulex* | 20,553 | 12 | 197,586 | 694 | 108.43 |
| *Drosophila melanogaster* | 24,420 | 10 | 172,853 | 1,019 | 169.90 |
| *Homo sapiens* | 252,301 | 12 | 30,347,128 | 90 | 77.11 |
| *Mus musculus* | 77,868 | 12 | 3,201,204 | 89 | 28.54 |
| *Thermus aquaticus* | 1,191 | 16 | 17,400 | 43 | 508.97 |
| Random sequence (A) | 3,441 | 12 | 227,399 | 217 | 34.41 |
| Random sequence (B) | 6,283 | 20 | 164,718 | 13,003 | 62.83 |
| Random sequence (C) | 24,498 | 12 | 40,229 | 83 | 244.98 |
| Random sequence (D) | 48,996 | 12 | 50,319 | 156 | 244.98 |
| Random sequence (E) | 154,142 | 12 | 7,414 | 24 | 1,541.42 |

***Tab. S13:*** *Fragment statistics for the in silico fragmentation with BsePI.*

| Species | No. fragments | Min. fragment length (bp) | Max. fragment length (bp) | Most freq. fragment length (bp) | Fragments per Mb |
|---|---|---|---|---|---|
| *Arabidopsis thaliana* | 1,111 | 8 | 962,949 | 8 | 9.28 |
| *Caenorhabditis elegans* | 8,340 | 8 | 226,241 | 8 | 83.16 |
| *Daphnia pulex* | 20,492 | 8 | 305,923 | 8 | 108.11 |
| *Drosophila melanogaster* | 15,997 | 7 | 229,453 | 8 | 111.30 |
| *Homo sapiens* | 80,301 | 8 | 30,545,250 | 8 | 24.54 |
| *Mus musculus* | 87,079 | 8 | 3,684,768 | 8 | 31.92 |
| *Thermus aquaticus* | 294 | 8 | 54,103 | 969 | 125.64 |
| Random sequence (A) | 3,415 | 8 | 201,945 | 8 | 34.15 |
| Random sequence (B) | 6,595 | 8 | 156,327 | 8 | 65.95 |
| Random sequence (C) | 24,269 | 8 | 47,822 | 8 | 242.69 |
| Random sequence (D) | 49,077 | 8 | 49,508 | 8 | 245.39 |
| Random sequence (E) | 154,252 | 8 | 8,378 | 8 | 1,542.52 |

***Tab. S14:*** *Complete list of the clusters of the selected peaks.*

→ see additional file ‘Supplement_RestrictionFragments_TabS14.xls’
